# Experimental evolution of *Chlamydomonas reinhardtii* in fluctuating versus constant temperature stress

**DOI:** 10.64898/2026.02.10.705100

**Authors:** Yeshoda Y. Harry-Paul, Rob W. Ness

## Abstract

Natural environmental variations are normal occurrences, however human-driven climatic changes continue to exacerbate the magnitude of these variations. This pushes organisms to their physiological limits where they must adapt or face extinction. The potential for organisms to adapt is of concern; however, the presence of fluctuating regimes to incorporate natural environmental variation is occasionally considered. Here, we investigate the standing variation in thermal optima of ten *Chlamydomonas reinhardtii* strains along a latitudinal gradient and determine whether adaptation to its temperature extremes is possible via long-term evolution. Despite their broad geographic origins, *C. reinhardtii* strains exhibited a similar thermal optimum (31.2°C - 32.4°C). We then evolved one strain for ∼900 generations under cold, hot and fluctuating growth conditions. Those grown in constant environmental conditions became specialized to their environment of 20°C or 37°C, whereas those evolved in a fluctuating regime performed comparably to the specialists, suggesting no long-term cost of adaptation via a fluctuating regime. We also found no differences in the amount of variation between fluctuating and constant conditions, suggesting similar amounts of divergence between conditions. Overall, despite our predictions that a fluctuating regime would hinder adaptation, we found that these lineages were able to adapt to this variable environment.

## Introduction

Climate change poses a significant threat to the stability and persistence of organisms and their associated ecosystems (Lancaster et al. 2022). As of 2015, there has been a 0.87°C increase in global mean surface temperature since the pre-industrial period of 1850-1900 (IPCC 2022). This human-induced global warming has already altered the climate in observable ways, including increased precipitation, sea level, land and ocean temperatures, as well as more frequent droughts and temperature fluctuations (e.g., heatwaves) (Salinas et al. 2019; IPCC 2022). The overall increase in temperature, coupled with additional heatwaves, can lead to extreme temperature exposure that organisms are not adapted to (IPCC 2022; Schaum et al. 2022; Schou et al. 2022). Organisms that are unable to mitigate temperature fluctuations via migration to favourable conditions often face extinction (Berg et al. 2010; Chevin et al. 2010) and thus must develop other adaptive mechanisms to cope with this range of temperatures (i.e. phenotypic plasticity or evolve genetic mechanisms) (Chevin and Hoffmann 2017; Diamond and Martin 2021; Sultan 2021). However, the interplay of these adaptive mechanisms and the use of one over the other in the adaptation to extreme temperature conditions is still under investigation, although both can be seen in literature (Harry-Paul et al. 2024; Roy et al. 2025).

Much of our understanding of how organisms adapt to tolerate extreme temperatures comes from experiments that measure performance at acute temperature shocks and across a range of temperatures to generate thermal performance curves (TPCs) (Ketola and Saarinen 2015; Ketola and Kristensen 2017; Schaum et al. 2022). The goal of TPCs is to provide an overarching view of temperature tolerance ranges and to predict how species could react to more extreme conditions. Generation of these curves generally involves subjecting organisms to multiple constant temperatures (Ketola and Saarinen 2015; Schou et al. 2017); however, natural environments are not constant and consist of natural fluctuations. In addition to natural environmental fluctuations, global warming can further amplify the variability in temperature that already exists (Schaum et al. 2022), leading to higher-than-normal temperature fluctuations. This can make predicting species’ future performance in extreme temperatures challenging (Schou et al. 2022), as studies have shown that thermal performance curves cannot be used to predict responses in a fluctuating environment (Ketola and Saarinen 2015). In addition, the period between environmental fluctuations has also been shown to affect the rate of adaptation, leading to more difficulties in predicting species survival (Ketola and Saarinen 2015; Schaum et al. 2022). As environmental fluctuations are common in nature, understanding how species and organisms respond to fluctuating or variable environments is crucial for understanding how species evolve in their natural habitats.

The adaptation to a fluctuating environment presents a distinct evolutionary and physiological challenge for many organisms. Contrary to stable environments, organisms must be able to tolerate a multitude of environmental conditions. How well organisms can adapt depends on the quality of the phenotype-environment match and whether the organism can change its phenotype to suit various environments (Levins 1968; West-Eberhard 2003; Diamond and Martin 2021; Sultan 2021). It is reasonable to expect that lineages that evolve in such a fluctuating environment will suffer trade-offs relative to those that evolve in a stable environment (Kassen and Bell 1998; Jacob et al. 2024). For example, if an environmental cue is required to predict an environmental change, there will likely be a cost to maintaining that cue recognition machinery (DeWitt et al. 1998; Schneider 2022). If the cue is unreliable or missed, this can create a mismatch between the organism and its environment, leading to a decrease in the quality of the phenotype-environment match (Schneider 2022). In addition, the adaptive mechanisms of the alternate environment must be maintained even in periods where that environment is absent, or risk a poor environmental match when the alternative environment returns (Levins 1968; Reboud and Bell 1997; DeWitt et al. 1998). Thus, the machinery and subsequent behaviour involved in the adaptation to more than one environment must be efficiently maintained, which is not required in a constant environment (Sultan and Spencer 2002). This increased energy expenditure on the maintenance of machinery for different environments can result in lower fitness within a given environment compared to a stable growth regime (Kassen and Bell 1998; Jacob et al. 2024). However, the cost of adaptation to a fluctuating regime remains a matter of controversy, as fitness benefits often mask costs associated with adaptation and thus must be studied depending on the biological context (Schneider 2022).

In addition, within a fluctuating environment, selection is no longer unidirectional and thus has less time to act on the genetic variation within each environment, weakening the strength of selection (Cvijović et al. 2015). As a result, slightly deleterious mutations which are beneficial in the alternative environment may be removed by selection at a slower rate, which leads to the conservation of genetic variation (Abdul-Rahman et al. 2021) and the silent accumulation of mutations (Kassen 2002; Lalejini et al. 2021). The presence of beneficial mutations linked to sub-optimal alleles which are beneficial in the alternative environment can result in generalist phenotypes with lower fitness in each environment relative to specialists, but higher mean fitness (Jacob et al. 2024). However, there is debate and conflicting evidence about whether adaptation to fluctuating conditions results in a generalist or specialist phenotype in one environment by sacrificing fitness in the alternative environment (Levins 1968; Kassen 2002; Räsänen et al. 2026). Within diatom communities, population fitness can increase as a result of the increase in fitness of one or two species (Wolf et al. 2024), similarly seen within adaptation to darkness in green algae (Bell and Reboud 1997). In these cases, the amount of variation within the population increases as individuals diverge in their phenotypic responses via the presence of evolved specialists and unadapted individuals. While both generalist and specialist evolutionary approaches can be seen in the literature, the conditions under which an approach is favoured are still an area that must be explored.

Evolution in a fluctuating environment may also influence the extent of phenotypic variance observed within and between populations. Environments that are heterogeneous or fluctuate over time can create dynamic fitness landscapes, which may select for distinct adaptive phenotypes that co-occur within populations or cause divergence among populations (Colegrave and Buckling 2005). Alternatively, the evolution of adaptive plasticity could allow phenotypic modifications in response to the environment, permitting lineages to occupy a single generalist fitness optimum and limiting phenotypic variance when measured in a single setting. Ultimately, the divergence of replicate lineages depends on the interplay between environmental predictability and the source of genetic variation. Populations with high standing variation may rapidly converge on similar optimal phenotypes, whereas populations that rely on de novo mutations to generate novelty are more likely to follow distinct evolutionary trajectories, resulting in greater phenotypic divergence (Barrett and Schluter 2008). We therefore expect that populations adapting to a fluctuating environment with limited genetic variation will generally increase in phenotypic variance relative to similar lineages evolving in stable environments.

Here, we use the unicellular green alga *Chlamydomonas reinhardtii*, which is a long-standing model for cell biology, photosynthesis and experimental evolution. As a member of the plant kingdom, it has many similarities to land plants, making findings generalizable to plants and other eukaryotes (Sasso et al. 2018; Fauser et al. 2022). In this study, we generate a broad-scale view of the natural variation in temperature tolerance among various natural genotypes originating from across a longitudinal gradient. In addition, while the physiology of acute temperature fluctuations has been investigated, no long-term experimental evolution to temperature has been conducted (Hemme et al. 2014; Kremer et al. 2018; Zhang et al. 2022). Our study consists of an investigation of genetic variation in temperature tolerance, followed by a long-term evolution experiment to cold, hot, and fluctuating temperature regimes, to answer the following questions: 1) Is there variation in the thermal optima across natural strains of *C. reinhardtii*? 2) Does constant long-term exposure to temperature stress evolve a temperature-adapted phenotype? 3) Does evolving under a fluctuating regime come at a fitness cost relative to lineages evolved in stable environments? and 4) Are fitness changes across replicate lineages more variable in fluctuating temperatures compared to constant growth regimes? In what follows, we present the fitness of 10 natural genotypes from along Eastern North America, at various temperatures, and show that thermal performance is remarkably similar despite a wide latitudinal range. Given that the minimal variation in T_opt_ between genotypes, rather than experimentally evolve multiple strains we used 12 lineages of a single strain were used to better understand the consistency of evolutionary responses without the confounding effects of variable genetic backgrounds. This design allows us to identify if adaptation to temperature stress can result in parallel evolutionary trajectories or are simply due to stochasstic events. Thus, we present the results of long term evolution of 48 replicate lineages adapted to cold, heat and a fluctuating temperature stress. We see that although all lines can adapt, the fluctuating regime adapted to a similar extent as the constant temperature lines within their specialized environments.

## Results

### What is the extent of variation in thermal tolerance across natural strains of C. reinhardtii?

To measure natural variation in temperature tolerance, 10 natural strains of *C. reinhardtii* along a latitudinal cline (Fig. 1) were grown in eight experimental temperatures ranging from 16°C to 39°C with a 25°C control (Fig. 2). Cells grown at cooler temperatures (≤ 20°C) did not achieve stationary phase within the time of the experiment due to slower growth (<u>Fig. S1</u>). However, cells grown at warmer temperatures displayed steeper exponential phases, indicating faster growth, although this effect disappeared at more extreme temperatures (≥35°C; <u>Fig. S2</u>). Cultures grown at extreme temperatures displayed a lower carrying capacity (<u>Fig. S1</u>) and underwent cell bleaching by the end of the growth trials. Indeed, we found a significant effect of temperature (F = 161.06, p < 0.001), cell strain (F = 19.8, p < 0.001) and their interaction (F = 3.48, p < 0.001) on growth. While the shape of the TPC for each strain is similar, the amplitude of the curves varies between strains, indicating differences in peak cell density at optimal growth temperature (Fig. 2). Despite these differences in cell density among strains, we found no evidence of a relationship between thermal optima (T_opt_) and latitude (Fig. 3). T_opt_ varied between 31.2°C and 32.4°C despite originating from locations as distant as Florida in the south and Quebec in the north. Among all 45 pairwise comparisons of T_opt_ ranges, 38% were significantly different between pairs (Fig. 3).

**Figure 1:**
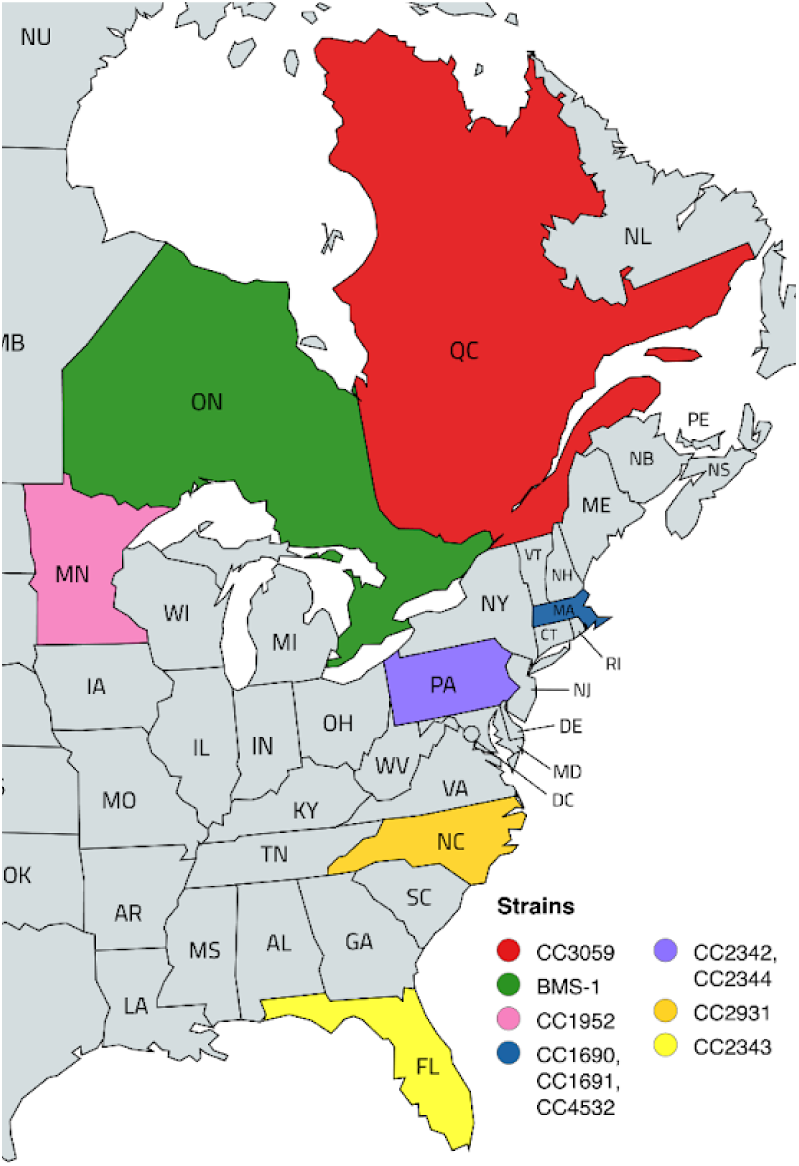
Locations of screened *C. reinhardtii* lines from Eastern Canada and the United States of America. Strains were selected to span the known latitudinal range in North America and to identify potential innate differences in thermal performance.

**Figure 2:**
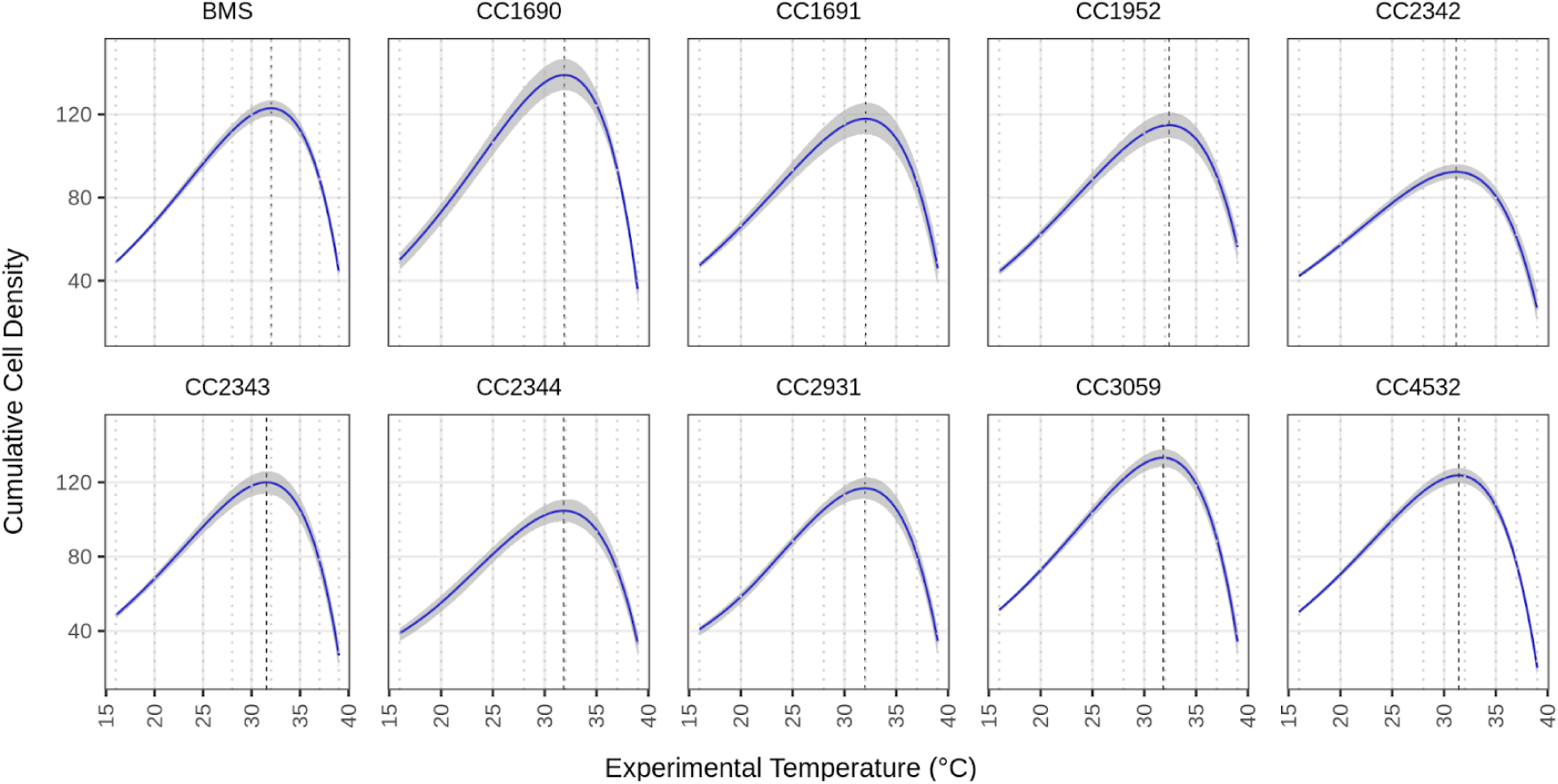
Growth of *C. reinhardtii* strains over various temperatures. Cumulative cell density is calculated as the area under the curve (AUC) of OD_680_ after four days of growth. The black dotted line indicates the optimal temperature. Grey dotted lines indicate experimental temperatures (16, 20, 25, 28, 30, 32, 35, 37, and 39°C).

**Figure 3:**
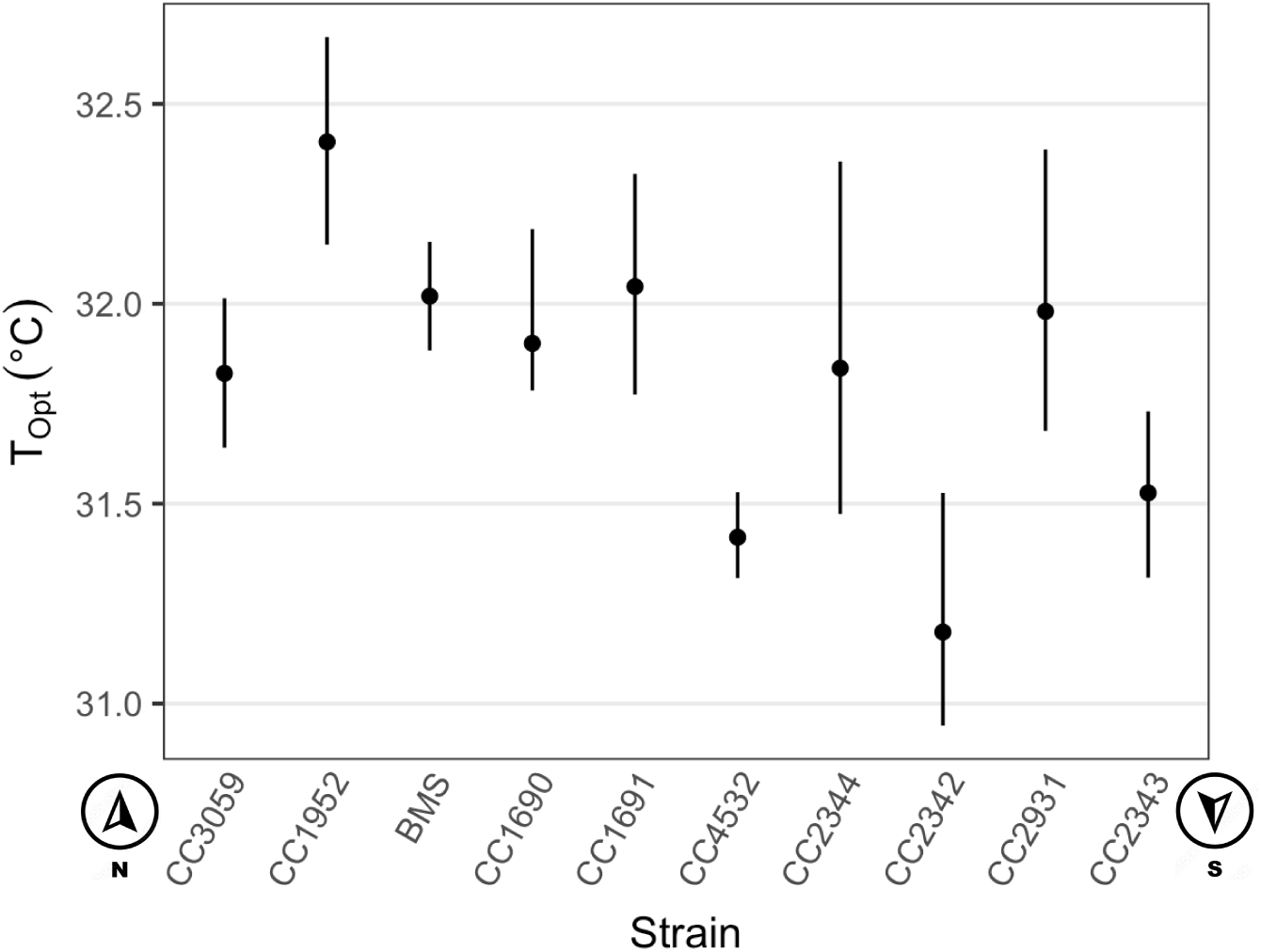
Optimal temperatures (T_opt_) for the 10 natural strains. Strains are arranged by latitude from north to south. Vertical bars indicate 95% confidence intervals.

### Does constant long-term exposure to temperature stress evolve a temperature-adapted phenotype?

We experimentally evolved 12 independent lineages in each of four temperature regimes (48 total lineages) to investigate adaptation to temperature stress. Specifically, 12 replicate lineages of the strain CC-4532 evolved in each of four experimental growth conditions: a 20°C constant (C20), 25°C constant (C25), which also served as the control, 37°C constant (C37) and a condition that fluctuated between 20°C and 37°C (Fluctuating). Hot and Cold experimental temperatures were chosen such that CC-4532 had similar amounts of growth (AUC) after 4 days in both temperatures (Fig. 2). These 48 lineages were propagated for nine months of experimental evolution (∼900 generations) within their respective conditions. The fitness of all 48 replicate lineages was assessed monthly in both 20°C and 37°C testing environments (Fig. 4).

**Figure 4:**
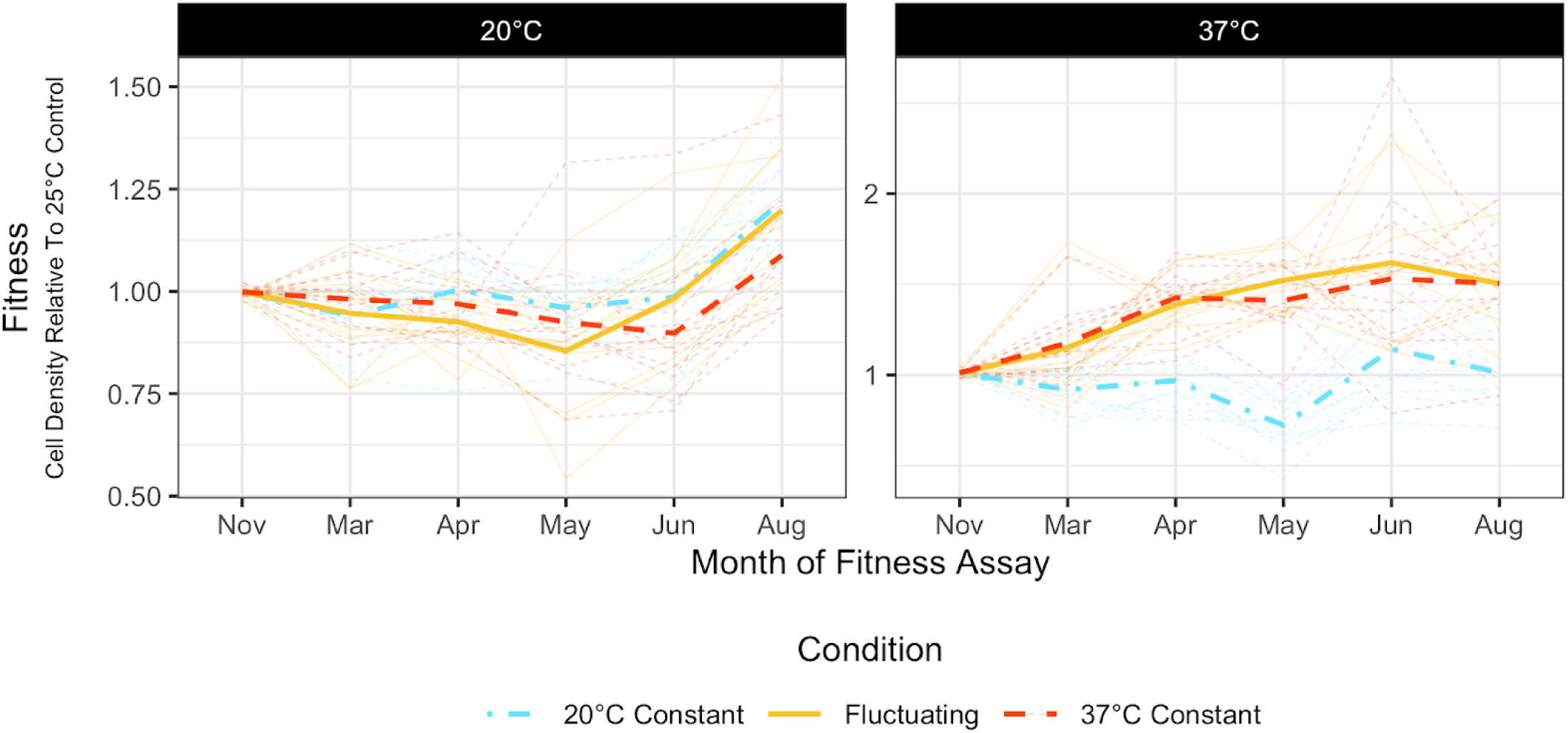
Fitness of the 12 experimental lineages within each condition measured in each experimental temperature over the course of the experiment. Fitness is reported relative to the 25°C control. Darker lines represent the average of the 12 lineages from each condition, while faint lines indicate each lineage. Solid lines, dashed lines and dot-dashed lines indicate the fluctuating, C37 and C20 lineages, respectively. Note that the y-axes are not conserved between plots.

To determine if lineages evolved to adapt to constant temperature regimes, we compared the fitness of both constant conditions relative to the 25°C control. Cumulative cell density was measured for each of the 12 lineages from C20 and C37 in both the 20°C and 37°C experimental environments. After nine months of evolution, a significant effect of Temperature (F = 19.1, p < 0.001), Condition (F = 17.4, p < 0.001) and their interaction (F = 89.7, p < 0.001) was found on cell density.

When measured in the 20°C environment, C20 lineages had significantly higher fitness compared to the control lineages with a 24% higher average fitness relative to the control (*t*(43.4) = -15.78, p < 0.001, Fig. 5). Although C37 lineages displayed lower fitness, this difference was not significant from C20 lineages (*t*(43.4) = 9.47, p = 0.128) or control lineages (*t*(43.4) = -6.31, p = 0.592). When measured in the 37°C environment, C37 lineages significantly outperformed the control, displaying a 51% increase in average fitness (*t*(43.4) = -27.95, p < 0.001, Fig. 5) and also significantly outperformed C20 lineages (*t*(43.4) = -27.69, p < 0.001). However, C20 lineages did not perform better than the control lineages (*t*(43.4) = -0.26, p = 1).

**Figure 5:**
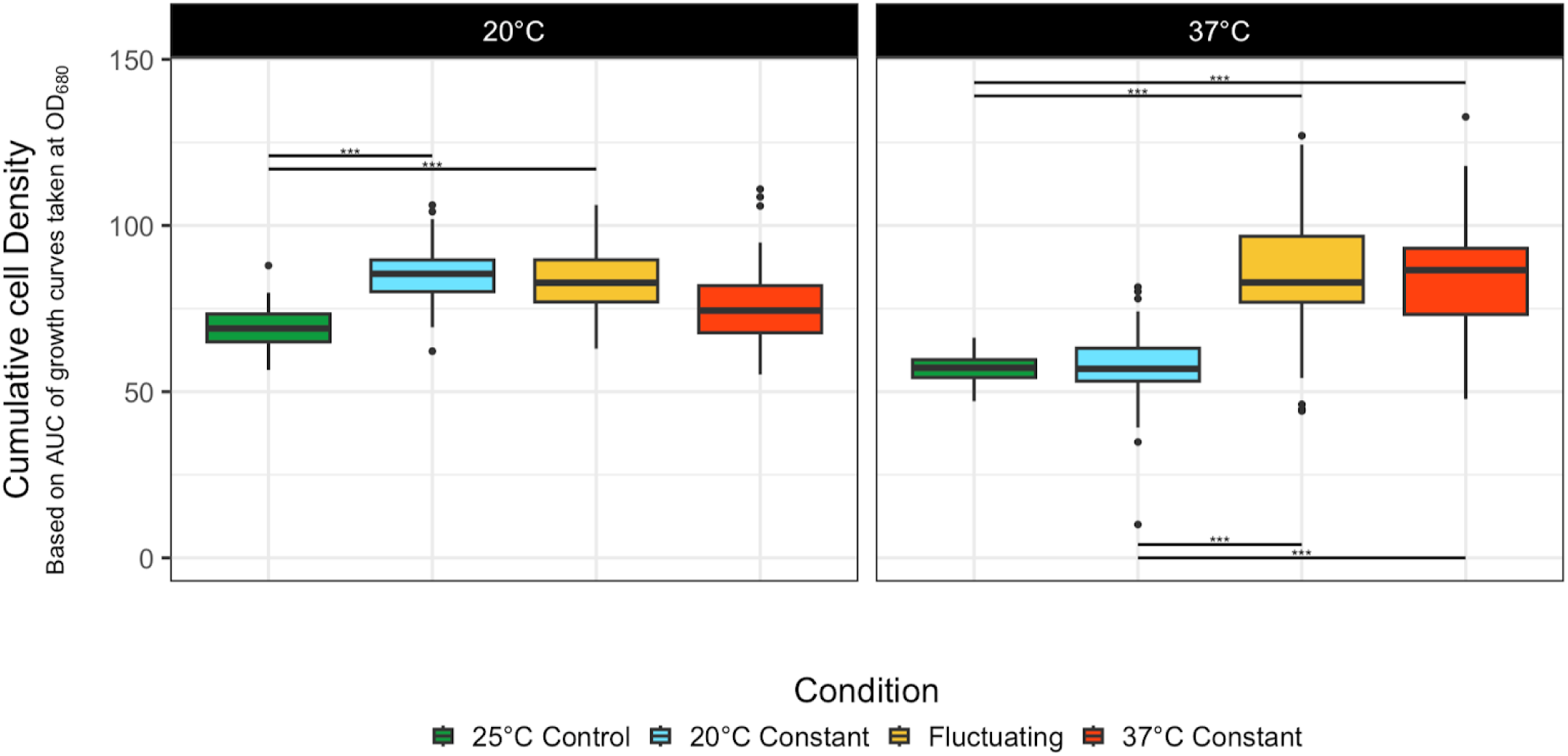
Final fitness of each condition measured in both 20 °C and 37 °C environments. *p < 0.05; **p < 0.01; ***p < 0.001.

### Does a fluctuating environment hinder the adaptation to temperature stress?

Above, we showed that the lineages evolved under constant conditions outperformed controls within their own specialized environments. However, we want to know if lineages evolved under a fluctuating regime will also develop tolerance levels similar to those of the specialists. Here, we compare the growth of the fluctuating lineages in both 20°C and 37°C environments with the growth of the respective specialist lineages within their environments and controls. Within the 20°C environment, lineages evolved under fluctuating conditions had significantly higher fitness than the control lineages (*t*(43.4) = -13.84, p = 0.0046); but did not significantly differ from C20 or C37 lineages (Fig. 5). Rather, the mean fitness of the fluctuating lineages fell between C20 and C37 lineages. Given that C20 lineages displayed the highest fitness within the 20°C environment, the similarity of the fluctuating lineages indicates comparable fitness to that of the 20°C constant lineages. Similarly, when assayed at 37°C, the fluctuating lineages had higher fitness than the control (*t*(43.4) = -27.99, p < 0.001) and C20 lineages (*t*(43.4) = -27.72, p < 0.001), but displayed comparable fitness to C37 lineages (*t*(43.4) = 0.04, p = 1, Fig. 5). Taken together, our data show that lineages evolved in a fluctuating environment were able to achieve fitness gains in 20°C and 37°C environments that were comparable to the lineages evolved under constant conditions, despite having spent approximately half of their evolutionary history in each condition.

### Is there more variance in fitness among lineages evolved in a fluctuating environment?

As each of the 12 replicate lineages within each condition evolved independently, their fitness may have diverged through the course of evolution. Indeed, we see a significant effect of the 12 replicate lineages in each of the evolutionary conditions (*X*^2^ = 107.02, p < 0.001) and in each experimental temperature (*X*^2^ = 20.14, p < 0.001). We also found significantly less variance among control lineages compared to the fluctuating and C37 lineages when fitness was measured in 20°C, and significantly less than all evolved treatments when measured at 37°C (Fig. 6). The significant effect of the 12 replicate lineages in each condition and temperature, as well as the increase in variance among lineages confirm that there was divergence among replicate lineages within the evolved lines. We also tested whether there was more variance between evolved lines from the fluctuating treatment compared to the two constant treatments. When fitness was measured in the 20°C environment, there was no significant difference in the amount of variance among replicate fluctuating lineages compared to either the C20 or C37 lineages (Fig. 6, <u>Table S1</u>). Similarly, at 20°C, there was no difference in the amount of variance in C20 or C37 lineages after Bonferroni corrections (F = 4.79, p_adj_ = 0.3669). Within the 37°C environment, there was a significant difference in the amount of variance between C20 and fluctuating lineages (F = 15.48, p_adj_ = 0.0017) as well as between C20 and C37 lineages (F = 10.73, p_adj_ = 0.0166). However, the amount of variance among the fluctuating lineages was not significantly higher than that of the C37 lineages (<u>Table S1</u>). Taken together, we see that although the replicates within each of the three evolutionary treatments increased in variance over the course of the experiment, the amount of variance among replicates from the fluctuating treatment was not consistently higher than the replicates from either constant temperature treatment.

**Figure 6:**
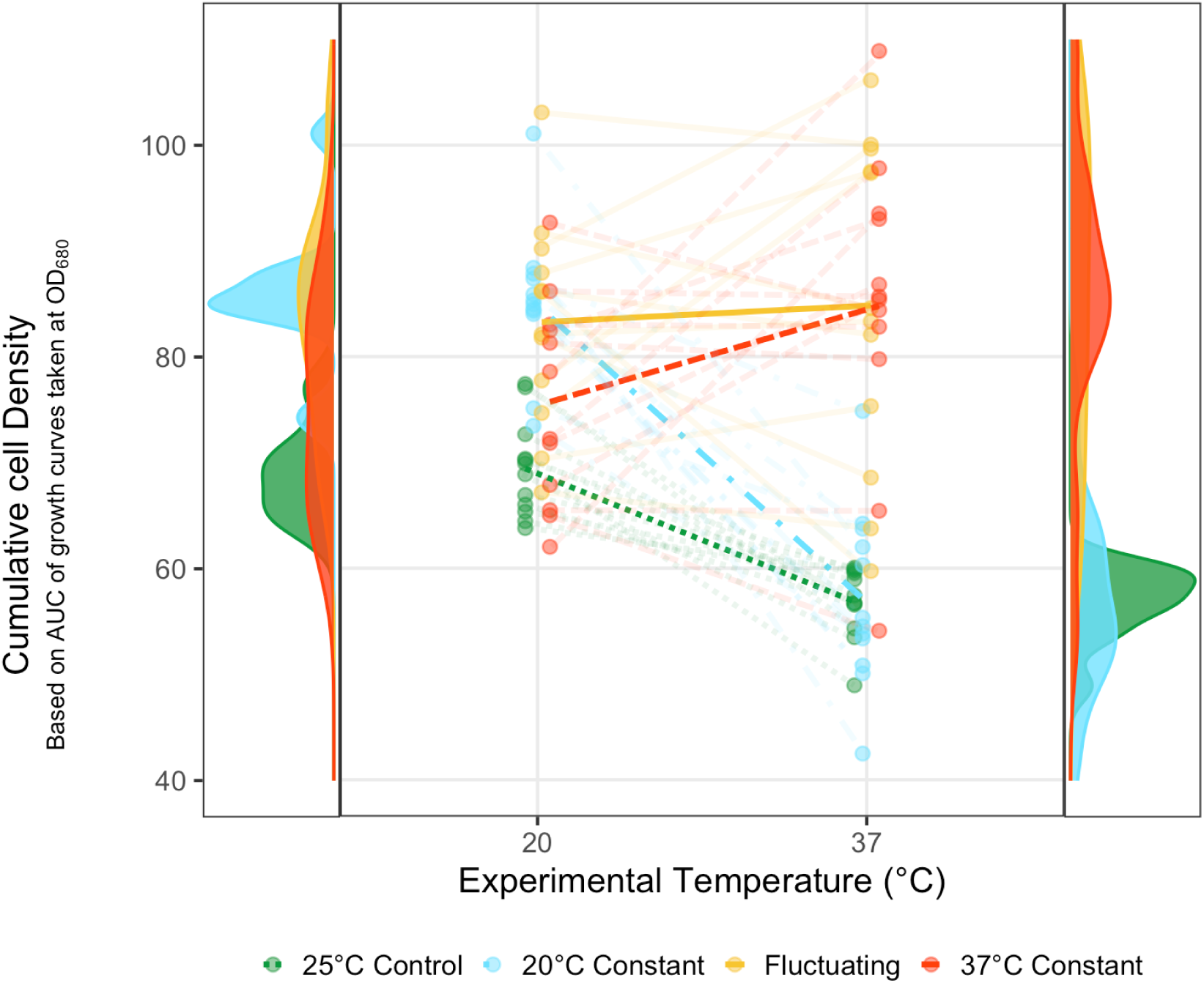
Distributions and mean fitness of each of 36 evolved lineages and control lineages, measured in both experimental temperatures. Mean of technical replicates for each of 48 independently evolved lineages (points), connected by faint lines to infer the relationship between environments. Dark lines connect the mean fitness of all 12 lineages between temperatures. Solid, dashed, dot-dashed and dotted lines denote fluctuating, C37, C20 and control lines, respectively.

## Discussion

To investigate the natural variation in thermal performance or T_opt_ of *C. reinhardtii*, we measured 10 natural isolates from across a latitudinal gradient in various temperatures (Fig. 2). Despite differences in origin and standard laboratory maintenance at 25°C, TPCs had a similar asymmetric shape, wherein growth was optimal at moderate temperatures but declined more steeply as temperature increased than as it decreased. Slower growth seen at cooler temperatures is seen in a variety of organisms (Mendoza and Cronan 1983; Wersebe et al. 2019; Ahmad et al. 2020), and is thought to be due to reductions in membrane fluidity and enzymatic constraints (Mendoza and Cronan 1983; Berry and Foegeding 1997; Nedwell 1999). Zhang et al (2022) showed that *C. reinhardtii* cultures subjected to acute heat shock (≥ 37°C) displayed increased chlorophyll levels but also compromised photosynthesis (Zhang et al. 2022). The initial increase in chlorophyll could result in rapid growth seen in our experiment; however, over time, this increase in chlorophyll, coupled with decreased photosynthesis, can increase the accumulation of reactive oxygen species (ROS) and lead to the degradation of chlorophyll and cell bleaching (Tchernov et al. 2004; Kobayashi et al. 2014; Buerger et al. 2020; Zhang et al. 2022). Similarly, increased membrane fluidity due to high temperatures can activate NADPH oxidase at the plasma membrane, also leading to increased levels of ROS (Königshofer et al. 2008). Further, due to initial rapid growth, nutrients may also be exhausted at a faster rate, resulting in limited nutrient availability to sustain the population in the stationary phase. In keeping with this, we see that temperatures over T_opt_ generally declined in total cell density over four days of growth, and in the extreme temperatures over 37°C, there was very little growth at all. The increased presence of ROS, compromised photosynthetic machinery, and lack of nutrient availability at increased temperatures create an environment of low fitness and provide an opportunity for evolution to occur.

While T_opt_ was statistically variable among the natural isolates, there was little biologically meaningful variation as the 10 strains’ optimum ranged from 31.2°C to 32.4°C (Fig. 3). Despite the drastic difference in mean surface temperature from Florida, USA, to Quebec, Canada, we saw no relationship between latitude or geographic origin and T_opt_. One possible explanation for the lack of differentiation could be that acclimation to laboratory conditions has removed locally adapted differences and canalized the thermal optima, as most of these strains have been in the lab for over 30 years (Craig et al. 2019). However, if the narrow range of T_opt_ is due to the selection for lab conditions, we would expect the T_opt_ to be closer to the standard lab growth temperature of 25°C instead of 31.8°C. Another possible explanation is that in nature, *C. reinhardtii* can lie as a dormant zygospore awaiting optimal conditions to germinate and begin the asexual part of its life cycle (Suzuki and Johnson 2002). Although northern latitudes, including Quebec, Ontario and Minnesota, are very cold in the winter, summer temperatures are high and overlap with temperatures experienced in Florida and North Carolina (refer to weather API from https://open-meteo.com). The small variation in T_opt_ may reflect the seasonal nature of *C. reinhardtii* growth and costs or constraints associated with altering growth temperatures. This relative lack of variation in the natural optimal temperatures and the fact that the optimal temperature does not match common laboratory conditions is even more indicative of such constraints, given the extent to which our experimental strains were able to adapt to higher temperatures.

Next, we investigated whether long-term exposure can lead to adaptation to stable or fluctuating temperatures. After nine months of evolution, lineages grown in constant temperature stress increased average fitness in their specialized environments by 24% and 51% for 20°C and 37°C, respectively (Fig. 5). The fact that the heat-stressed lineages increased their fitness more than twice as much as the cold-adapted lineages is noteworthy. This could be explained by stronger selective pressure in the 37°C environment relative to the 20°C environment. While we showed that net growth in 20°C and 37°C was similar in the ancestor (Fig 2), the 37°C environment prompted an earlier exponential phase that did not reach high carrying capacity due to cell death. In contrast, the 20°C environment resulted in a delayed exponential phase but less cell death (<u>Fig. S2</u>). As a result, while the cumulative cell density after four days of growth is similar, the increased response to selection in the 37°C environment could have resulted from the higher cell division rate of C37 lineages and more total generations, allowing for more selection to occur within the same time frame. Another explanation for the increased fitness observed in C37 lineages is that there may be more, or stronger effect beneficial mutations available. This is of particular importance in our experiment, as isogenic lineages must be dependent on *de novo* mutations for all genetic variation. Under native lab conditions, the mutation rate of *C. reinhardtii* is ∼1.2 x 10^-9^ (Ness et al. 2015), thus we would expect ∼58 mutations in each lineage under neutrality. We do not know how temperature stress affects mutation rate in *C. reinhardtii,* but from other studies done in *E. coli* (Chu et al. 2018), *C. riparius* (Waldvogel and Pfenninger 2021) and *D. melanogaster* (Muller 1928), there is reason to think that both heat and cold stress can increase the mutation rate. We also know from heat shock response studies of *C. reinhardtii* that there is a large complex of responses that are triggered when cells experience high temperatures (Hemme et al. 2014; Magni et al. 2018; Zhang et al. 2022). This plastic genetic mechanism could provide a potential source of evolutionary novelty to be canalized within a high-temperature-adapted lineage (Waddington 1953; Crispo 2007). While there is also a large complex of responses in reaction to cold stress (Valledor et al. 2013; Li et al. 2020; Peng et al. 2021; Abeysinghe et al. 2022), many of the large-scale responses occur at temperatures below what has been studied here (e.g. 5°C). Thus, 20°C might not offer a strong enough pressure to elicit a large cold-stress plastic response, nor allow enough generations (Valledor et al. 2013) to canalize the response. The higher division rate in heat, in tandem with a higher mutation rate, provides an increase in genetic material and time available for selection and thus an increase in the rate of evolutionary adaptation.

It is often predicted that rapid adaptation to a strong selective pressure will result in the fixation of large-effect mutations that may be harmful in the ancestral or other environments, resulting in lower fitness (Mackay 2001). This trade-off could be the result of antagonistic pleiotropy associated with mutations that underpin each specialized phenotype, leading to higher fitness in one environment with a regression in the alternate environment (Levins 1968; Kassen and Bell 1998; Diamond and Martin 2021). However, while we expected that adaptation to one environment would result in a reduction in fitness in the alternate environment, this was not the case (Fig. 5). Rather, C20 and C37 retained the ancestral fitness of the alternate condition, as evidenced by the non-significant differences between each of the constant conditions and the control within the alternate environments (Fig. 5).

We similarly saw relatively little evidence for a trade-off of adaptation to the two temperatures in the fluctuating lineages. Early in the experiment, lineages in the fluctuating growth regime adapted quickly to the 37°C environment, with no significant adaptation to the 20°C environment (Fig. 4). This asymmetry could reflect the same set of differences we outlined above when interpreting the difference in adaptation between C20 and C37 lineages. However, we predicted that the fluctuating lineages may experience higher selective pressure in the 37°C environment, which could result in a trade-off that reduced performance in the 20°C environment (Cheng et al. 2024; Wolf et al. 2024; Roy et al. 2025). However, over the latter half of the experiment, the fluctuating lineages began to adapt to the 20°C environment and subsequently, by the end of the experiment, the fluctuating lineages adapted to both environments such that they were comparable to both the C20 in 20°C and C37 in the 37°C environment (Fig. 5). Many evolutionary theories assume that the area under fitness curves are conserved, such that adaptation to temperature fluctuations is expected to produce generalist phenotypes with a wide breadth of temperature tolerance at a reduced fitness compared to specialists (Levins 1968; Gilchrist 1995; Ketola et al. 2014). Indeed, previous studies have shown that fluctuating growth regimes performed worse than constant conditions when measured at constant temperatures (Ketola and Saarinen 2015; Jacob et al. 2024), suggesting different mechanisms that potentially compete for shared resources (Saarinen et al. 2018). However, we found that there was no significant difference in the performance of the fluctuating lineages compared to the constant lineages when tested within constant conditions (Fig. 5), suggesting the area under fitness curves is not necessarily conserved. Despite less time for selection to act, this pattern of equal or better performance in fluctuating regimes compared to specialists reared at constant conditions is seen elsewhere (Reboud and Bell 1997; Kassen and Bell 1998; Ketola et al. 2013; Fasanello et al. 2024; Wolf et al. 2024). However, we see markedly different results than a recent study that adapted fission yeast to constant and fluctuating temperature, which found that constant temperature was a stronger selective pressure than variable temperatures (Räsänen et al. 2026). While it is difficult to know why the results differ, the fact that *C. reinhardtii* needs to acclimate to a wide variety of temperatures in nature may predispose it to variable temperature adaptation.

In addition, the time between fluctuations also plays a role in understanding the rate of adaptation to novel environments. Theory predicts that fine-grained environments, where individuals experience multiple different environments within a generation, can elicit a reversible phenotype via phenotypic plasticity, resulting in more of a generalist phenotype compared to fluctuations that scale over generations (Kassen and Bell 1998; Kassen 2002). While initial adaptation to a fluctuating regime can depend on time between fluctuations as well as acclimation history (Kremer et al. 2018), the end adaptive result is the same (Schaum et al. 2022). Thus, while fluctuations of 4-5 days facilitated adaptation to both environments after ∼900 generations, a shorter fluctuation time may have resulted in a faster rate of adaptation, but adaptation nonetheless.

Under strong selection, independent evolutionary lineages may acquire different de novo mutations, potentially increasing variance among those lineages over time (Bell and Reboud 1997; Goho and Bell 2000; Colegrave and Collins 2008; López-Cortegano et al. 2021). We indeed observed that when variance among replicate lineages was measured from each of the evolutionary treatments, there was an increase in the amount of variance relative to the control lines that were maintained under standard lab conditions (Fig. 6, <u>Table S1</u>). Given that all lineages were evolving for similar amounts of time, this supports the prediction that directional selection has driven divergence among lines, even those subject to the same selection pressure. Given that each lineage was isogenic, the lack of gene flow and sexual reproduction, all adaptive mutations must have arisen from de novo mutations (Barrett and Schluter 2008). As a result, the rate and distribution of fitness effects must have been the primary driver of the amount of variation among lines. We observed that relative to the mean fitness, chance variance among lineages was high, which suggests a low rate of beneficial mutations or highly variable fitness effects of those mutations. We also expected increased variance in the fluctuating lineages, as given their heterogeneous environment, they may accumulate mutations which benefitted one temperature environment over the other. However, this hypothesis was not supported by the data, as the amount of variance among fluctuating lineages was not significantly different from that of the other evolved lines. A more prominent trend was a 15.8- and 18.3-fold increase in variance among C37 and fluctuating lineages than controls when grown at 37 °C. This is in contrast to Ketola et al (2013), where lineages in fluctuating selection environments accumulated less variance than constant temperature lines (Ketola et al. 2013). However, our experiment was significantly longer, and thus there was more time for lineages to accumulate fitness changes and diverge from each other. It seems likely that in the long term, replicate lines will ultimately converge on a similar fitness level, and variance will diminish. Over the time frame of the current study, lineages have not yet reached that state.

In conclusion, we identified that while there is variation in T_opt_ across a latitudinal cline of *C. reinhardtii*, this variation only spans 1.2°C (Fig. 3). Whether this is due to the seasonal growth patterns of *C. reinhardtii* or to acclimation to lab conditions is unclear. In addition, after ∼900 generations of evolution within 20°C and 37°C environments, lineages subjected to constant conditions performed the best within their specialized environments (Fig. 5). However, the fluctuating regime was not significantly different from either condition within its specialized environments, suggesting that a fluctuating regime can perform just as well as the constant conditions. In addition, we did not find a significant difference in the amount of variation between fluctuating and constant reared lineages within their specialized temperature (Fig. 6). Therefore, we’ve displayed here the adaptive potential for *C. reinhardtii* to adapt to novel environments, suggesting the capacity to survive in an ever-changing climate.

## Methods

### General Growth conditions

Strains were cultivated in liquid Tris-Acetate-Phosphate (TAP) media with continuous light and shaking for four days. Cell cultures were grown in continuous light at ∼6000 lux with continuous agitation (100 RPM) in an Infors HT multitron standard incubator.

### Thermal Performance Curves

Cell culture was inoculated into 1 mL of Tris-Acetate Phosphate (TAP) media from a 1.5% TAP-agar slant into a 24-well plate using a 1 μL inoculating loop. After four days of growth, 11 μL of culture was spread on a 1.5% TAP-agar dish. After three days, eight colonies were picked, representing eight biological replicates, and inoculated into 1 mL of TAP in a 48-well plate. After four days of growth at 25°C, the culture was diluted 1:50 into a 24-well plate. Three days later, approximately 20,000 cells (via Beckman Coulter Cytoflex) were inoculated in 200 μL of TAP in a 96-well plate and grown at 16, 20, 28, 30, 32, 35, 37, and 39°C with a corresponding 25°C control. OD_680_ (BioTek Epoch reader) was used as a measure of total chlorophyll content to construct growth curves, from which the area under the curves was calculated to infer cell density (Kholssi et al. 2023). While OD_680_ was reported here, the findings are similar to those with OD_750_ (<u>Fig. S3</u>).

### Evolution Experiment

To create the initial ancestral population, a 1 μL loop of CC-4532 from a 1.5% TAP-agar slant was inoculated into 1 mL of TAP. After three days, the culture was streaked onto a 1.5% TAP-agar plate from which the 12 original biological replicates were picked. Here, 12 lineages of a single strain were used to better understand the consistency of evolutionary responses and identify if adaptation results in parallel evolutionary trajectories. These colonies were inoculated into 1 mL of TAP in a 24-well plate. Culture was diluted 1:15 in 1.5 mL of TAP and left to grow for five days at 25°C. The regrowth phase ensured robust growth of the cell culture. Four replicate 24-well plates were made, each with a cell concentration of 300-500 cells/μL. Cell concentration was measured via flow cytometry (Beckman Coulter Cytoflex), using FSC. Each plate was placed in the temperature associated with its respective condition (control at 25°C, 20°C specialist at 20°C, 37 °C specialist at 37°C and the fluctuating beginning at 37°C). Cultures were transferred into fresh media every 4-5 days. Each transfer day, the fluctuating regime was placed in the alternate temperature environment.

### Fitness Assay

To assess the adaptation of each condition to its respective temperature, fitness assays were done every 4-6 weeks during the evolution experiment. A replicate plate of each condition was made for every fitness assay, such that the original culture was not disturbed during the experiment. The replicate plates were placed into a 25°C environment for 4 days to acclimate to a neutral condition and prevent the carryover of previous environmental effects (Wolf et al. 2024). Each plate was then split into two equal-volume replicates, diluted 1:1 in fresh TAP media, and placed into either the 20°C or 37°C environment to precondition the cultures. In this way, cultures are exposed to the same temperature and have time to adapt to their testing condition before their fitness is measured to minimize the carry-over of any adaptive advantage from their original environments. After one day of growth, approximately 20,000 cells were used to inoculate five replicates of each culture from each condition into 96-well plates. Replicates were measured at 20 and 37°C for four days and assessed twice daily via spectrophotometry at 680nm (BioTek Epoch reader). Cell concentration was measured on initial inoculation days and day four, via flow cytometry (Beckman Coulter Cytoflex).

### Statistical Analysis

#### Temperature Screen

With R v4.4.3(R Core Team 2025), we used the function `lmer` (*lme4* v1.1.3)(Bates et al. 2015) to fit linear mixed-effect models for these analyses. To investigate the presence of batch effects, cell line and replicate were modelled as random effects with cell density as the response. Cell density was calculated as the cumulative area under the growth curve for each time point using the function `area_under_curve` (*bayestestR* v0.17.0(Makowski et al. 2019)) with the spline method. The full model used in the remainder of the analyses was as follows: Cell Density ∼ Temperature * Cell Line + (1|Batch) + (1|Replicate), based on a lowest AIC score. The function `anova` was then used to generate summary statistics.

Models were fit to randomly sampled cell densities with replacement at each temperature within each cell line. Thermal performance curves were fitted using the *rTPC* v1.0.4 (Padfield et al. 2021, 2025) package with the rezende_2019 (Rezende and Bozinovic 2019) model chosen *a priori*. While a quadratic model resulted in a low AIC, it did not follow the expected thermal performance curve shape as mentioned in other literature (Kellermann et al. 2019), and thus, the Rezende model was chosen. This was repeated 10000 times for each cell line to generate 95% confidence intervals of the T_opt_.

#### Fitness Assay

Cell density was calculated using the same area under the curve method as the temperature screen. The cell density for the 20°C, 37°C and fluctuating conditions was normalized via per colony division of the median cell density of the 25°C control. There was no effect of replicate, and thus it was represented as a random effect. The following linear model was used for the remainder of the analysis: Area ∼ Condition Environment + (1|Colony) + (1|Colony:Condition) + (1|Colony:Environment), with the `anova` and `ranova` functions to generate summary statistics. Pairwise comparisons were made using the `emmeans` function (*emmeans* v 1.11.2)(Searle et al. 1980).

#### Among line variance comparison

Comparisons of variance among lines within each evolution condition were done via subsetting the data into pairs of conditions and running a Levene test via the `leveneTest` function (*car* v3.1.3)(Fox et al. 2024) for each environment. P-values were adjusted via Bonferroni corrections using the `p.adjust` function (stats v4.4.3). (R Core Team 2025)

## Acknowledgements

The authors wish to thank Stephen Wright, Keiko Yoshioka and Ru Zhang for helpful insights and suggestions, as well as Adrian Nuredini, Kyli Kindree, Ayesha Syeda, and Maja Tingchaleun. This work was supported by a Natural Sciences and Engineering Research Council Discovery grant to RWN (RGPIN/06331-2016) and an NSERC graduate scholarship to Y.Y.H.P.

