## Supplementary material for "Experimental evolution of *Chlamydomonas reinhardtii* in fluctuating versus constant temperature stress": Fig. S1;Fig. S2;Table S1;

### ***Supplemental***

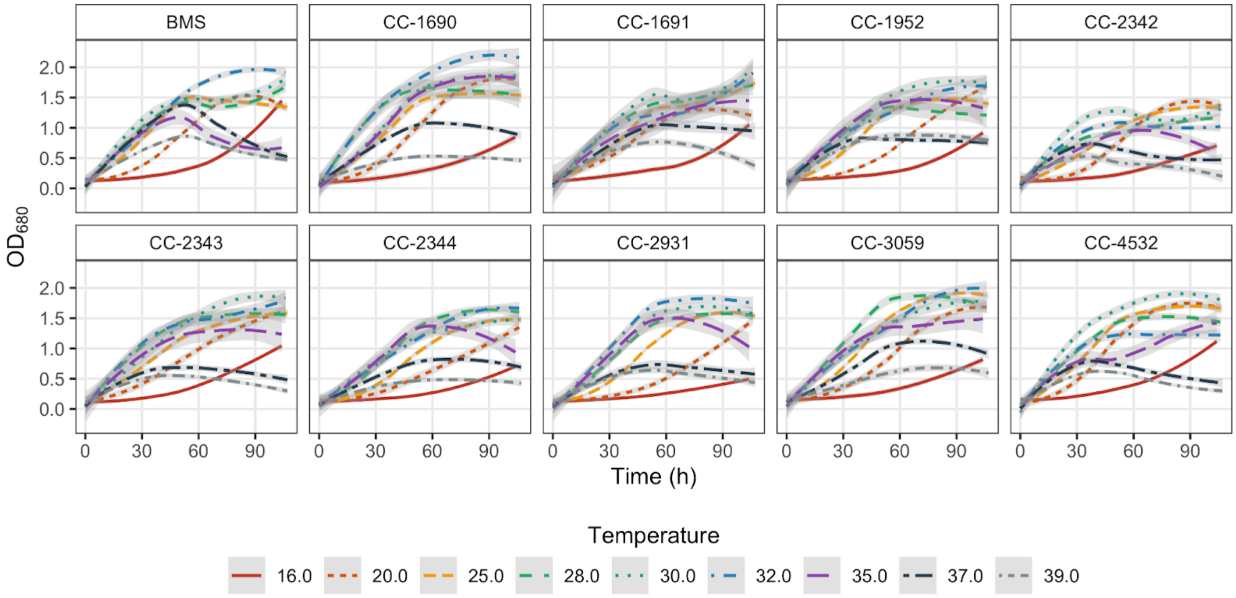


Figure S1: Raw growth curves of *C. reinhardtii* strains over various temperatures. Shaded regions correspond to the standard error of the curves.

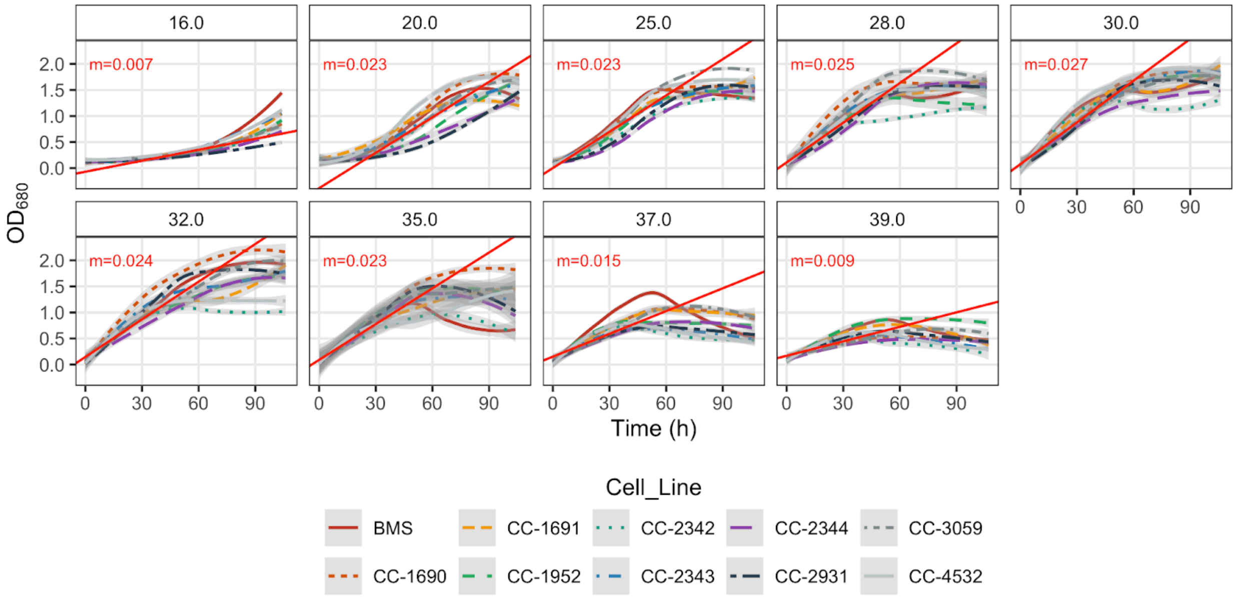


Figure S2: Raw growth curves of *C. reinhardtii* strains over various temperatures. Shaded regions correspond to the standard error of the curves. The red line indicates the linear trendline of the exponential growth phase.

Table S1: P-values before and after Bonferroni corrections for Levene's test on the amount of variation between conditions within the respective temperatures

| Temperature | Pairs | F | Raw p-values | Adjusted p-values |
|---|---|---|---|---|
| 20 | Fluctuating, C37 | 0.47 | 4.932120e-01 | 1.0000 |
| 20 | Fluctuating, C20 | 2.70 | 1.030873e-01 | 1.0000 |
| 20 | C20, C37 | 4.79 | 3.057467e-02 | 0.3669 |
| 20 | Fluctuating, C25 | 13.83 | 3.078764e-04 | 0.0037 |
| 20 | C20, C25 | 4.89 | 2.898548e-02 | 0.3478 |
| 20 | C37, C25 | 15.91 | 1.157347e-04 | 0.0014 |
| 37 | Fluctuating, C37 | 0.42 | 5.187749e-01 | 1.0000 |
| 37 | Fluctuating, C20 | 15.48 | 1.411887e-04 | 0.0017 |
| 37 | C20, C37 | 10.73 | 1.382285e-03 | 0.0166 |
| 37 | Fluctuating, C25 | 51.26 | 7.451062e-11 | 0.0000 |
| 37 | C20, C25 | 12.92 | 4.745122e-04 | 0.0057 |
| 37 | C37, C25 | 42.47 | 1.848332e-09 | 0.0000 |

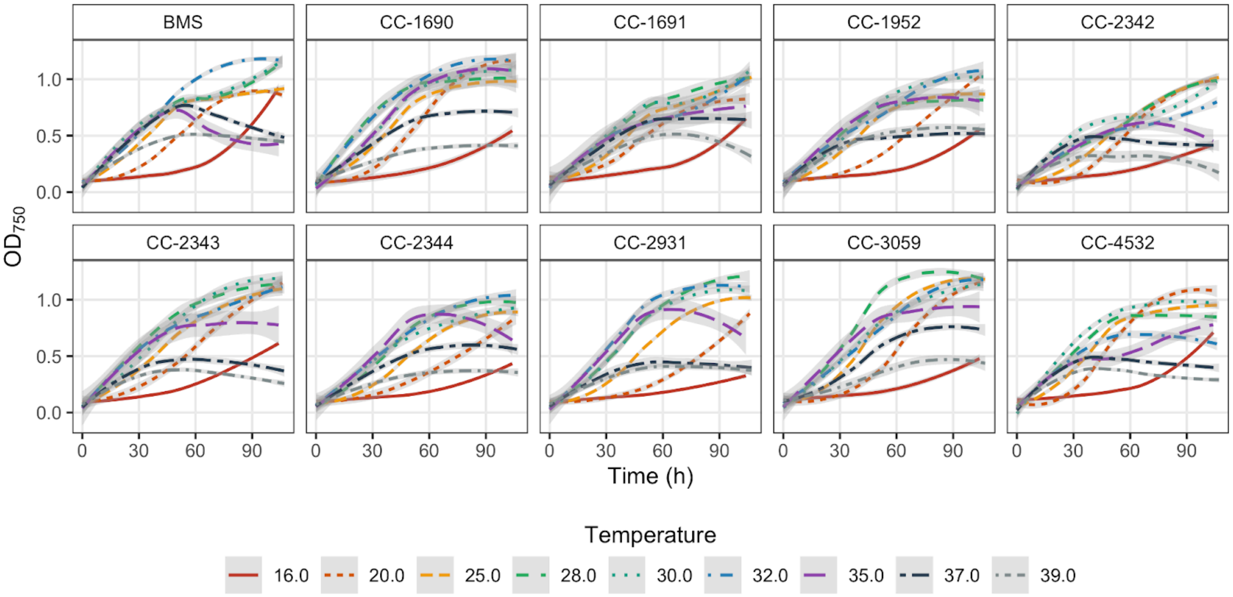


Figure S3: Raw growth curves of *C. reinhardtii* strains over various temperatures at $OD_{750}$. Shaded regions correspond to the standard error of the curves.
